# Development of a quantitative colorimetric LAMP assay for fast and targeted molecular detection of the invasive lionfish *Pterois miles* from environmental DNA

**DOI:** 10.1101/2023.12.20.572323

**Authors:** Katherine Hartle-Mougiou, Chrysoula Gubili, Panagiota Xanthopoulou, Panagiotis Kasapidis, Martha Valiadi, Electra Gizeli

## Abstract

The Mediterranean basin has seen an increased influx of invasive species since the Suez Canal expansion in 2015. The invasive lionfish species, *Pterois miles*, has rapidly established new populations in the Eastern Mediterranean Sea, impacting local fish biodiversity. Here, we have developed a new, fast (< 35 min) molecular approach to detect and quantify *P. miles* environmental DNA (eDNA) in combination with a portable device for field-based analysis. Using a species-specific real-time colorimetric loop-mediated isothermal amplification (qcLAMP) for the cytochrome oxidase subunit 1 (COI) gene, we demonstrate a high sensitivity with a limit of detection of 0.002 ng DNA per reaction, equivalent to only 50 copies of the COI gene. The assay is specific to the target in the presence of closely related and co-occurring species, and it is quantitative over five orders of magnitude. We validated the assay using aquarium water samples and further demonstrated its utility on natural eDNA samples collected from locations around the island of Crete where *P. miles* had been sighted. *P. miles* was indeed detected in three out of nine locations, two nature reserves and a closed bay. Lack of detection in the remaining locations suggest that populations are still at a low density. We also demonstrate the feasibility of *P. miles* eDNA qualitative detection directly from the filter used to collect eDNA-containing particles, completely omitting DNA extraction. Overall, we present a new approach for fast and targeted eDNA quantification. The developed LAMP assay together with the quantitative real-time colorimetric detection approach open new possibilities for monitoring invasive *P. miles* in the field.

## Introduction

Marine biological invasions are increasing globally, driven mostly by human-mediated transportation and elimination of natural barriers to dispersal, and further facilitated by global warming (1). Invasive species damage ecosystem equilibria in their new habitat, by causing new ecological pressures and benefitting from a lack of native predators (2). These invasions have detrimental socio-economic impacts (3), and some even pose a threat to human health (2). The Mediterranean Sea is one of the most vulnerable regions for biological invasions, 993 non indigenous species were reported by the end of 2021, with 751 species confirmed as established (4). Two key environmental barriers between the Red Sea and Mediterranean Sea have been removed, facilitating species migration. Continued expansions of the Suez Canal have removed the channel’s hypersaline regions (5) whilst the eastern Mediterranean Waters have rapidly warmed to similar temperatures as the Red Sea (6, 7).

One of the most significant ongoing invasions in the eastern Mediterranean Sea is by the Indo-Pacific lionfish species, *Pterois miles* (Bennett, 1828), a fish with long venomous spines of the Family *Scorpaenidae*. Its indistinguishable relative, *Pterois volitans*, have already caused significant damage in the North West Atlantic since the 1980, causing decline in native fish species and affecting reef communities with associated economic impacts (8). Their success is due to their high fecundity and spawning activity, lack of natural predators in invaded regions, their poisonous fins, broad diet, and ability to occupy various habitats and depths (9-11). The invasion in the eastern Mediterranean is at earlier stages, with reports of established populations commencing in 2015 (12). In the past few years, populations have primarily affected the coasts of Israel, Syria, Lebanon, Turkey and Cyprus and more recently Greece (11). Despite large-scale efforts by the EU H2020 RELIONMED programme to collect data in order to officially include this species in the EU Invasive Alien Species list and implement management strategies to intercept the invasion, this has yet to happen (13). Management strategies have already been tested in the Atlantic and recommendations have been made for population control by targeted removal and commercial exploitation (14). The Mediterranean lionfish invasion front is currently situated in Greece with populations rapidly establishing in the Levantine Sea, Aegean and Ionian Seas; further spread is limited by the 15.3 °C isotherm to the north and west although there have been a few isolated sightings in Sicily and Croatia (8, 15, 16). The Island of Crete, in Southern Greece, has seen increasing populations in the last five years (12), with lionfish now regularly available at fishmongers.

Early intervention on marine invasions requires early detection of alien species, as well as information about population abundance and habitat preferences. Current methods for marine organism observation include visual surveys (snorkelling and SCUBA diving), observations by citizens, and trawl nets (7, 17, 18). However, these approaches are ineffective for low density populations as, when individuals are infrequent, they can be easily missed by divers (19). Conversely, molecular detection of species from environmental DNA (eDNA) is more sensitive and cost efficient for low density populations at the initial phase of an invasion (19). eDNA is genetic material contained in free particles that have been shed from organisms in the ocean e.g., from urine, faeces, gametes, and skin (20). Species-specific eDNA-based approaches (21) have been demonstrated for monitoring invasive fish such as carp and brook trout fish in USA freshwaters (20, 22, 23). On the other hand, eDNA metabarcoding, which profiles eDNA from the whole community, might miss rare species due to insufficient sequencing depth or lack of sequences on the reference database. Additionally, this approach requires sending the sample to a dedicated laboratory and time-consuming bioinformatic analysis, increasing the cost and analysis time (24).

Species-specific molecular detection is the preferred approach for early detection and spatial tracking of specific species of concern. Quantitative PCR (qPCR) and digital droplet PCR (ddPCR) have been used in numerous quantitative eDNA studies as they have high sensitivity in detecting eDNA targets and provide quantitative estimations (25-30). DdPCR is less prone to errors and more accurate than qPCR, especially at low eDNA concentrations, making it better suited to provide abundance and biomass estimates of the target species (28). Assays need to be carefully optimized to ensure target specificity, as eDNA samples contain genetic material from various organisms, including species that are closely related to the target organism. Quantitative data generated by these approaches enable estimates of population size, helping to identify preferred habitats, potential breeding grounds, and define modelling approaches that account for currents and life cycle history in eDNA based ecosystem studies (19, 28, 31-33). However, both aforementioned PCR-based methods are lab-based, expensive (including both the assay reagents and instrument) and require purified DNA samples.

Given the need for intensive field surveys, there will be significant benefit from approaches that provide results quickly and at low cost. Ideally, these tests should be performed on site, to avoid the need to transport sensitive eDNA samples to a laboratory freezer in strict conditions. This approach would enable field studies with improved geographic coverage especially at remote locations. To this end, the characteristics of isothermal nucleic acid amplification enable analysis in small, simple and portable devices, as they obviate the need for energy-demanding thermal cycling as in PCR (26, 34). Isothermal amplification methods amplify DNA based on strand displacement enzymatic activity, instead of temperature cycling. For marine environments with high biodiversity, loop-mediated isothermal amplification (LAMP) stands out as a method of choice for eDNA analysis due to its high specificity. This is due to the requirement of 4-6 primers to be used to recognise 6-8 regions of the target. LAMP also has a very high sensitivity due to high product yield and is a very robust method even in the presence of complex samples (34). Detection of LAMP amplicons can be achieved in real-time to enable target quantification by including a fluorescent DNA dye; this method is primarily lab-based and relatively expensive. An alternative approach includes the use of pH sensitive dyes and visual assessment of colour change. However, although of low cost and suitable for field testing, visual assessment suffers from low sensitivity while it is also subjected to the user’s interpretation and requires a “trained eye” for accurate monitoring (35).

Few portable systems have been developed for marine applications so far, with the main focus being on the detection of harmful algae (36). These are mainly based on assays targeting ribosomal genes which exist in hundreds of copies per cell to enhance the assay sensitivity. These assays are qualitative (target presence / absence only) due to a lack of portable instrumentation with enough accuracy to enable target quantification. Nevertheless, LAMP assays have been shown to achieve a high sensitivity which is ideal for marine eDNA studies; although this is reported in non-uniform units, the Limit of Detection (LoD) is in the range of 1 cell mL^-1^ for LAMP with a fluorometric dye and an instrument measuring turbidity in the reaction (37) and 0.034 pg DNA per reaction with detection using lateral flow strips (38). In our lab, we have developed a dedicated portable device to improve colorimetric measurements and enable quantitative analysis of the samples. The device can record the colour change in the LAMP reaction-tube via an embedded mini-camera; the device has been demonstrated to be suitable for fast and sensitive testing of swab samples for SARS-CoV-2 RNA and cancer genetic mutations in patients’ tissue (39). Overall, the method has been adapted to provide real-time results with reduced complexity and user-friendly smartphone operation, which can be used in the field.

In this study, we aimed to develop a real-time colorimetric loop-mediated isothermal amplification (qcLAMP) assay for targeted estimates of *P. miles* presence and population density through eDNA. This assay is implemented with the newly developed portable device to enable readiness as an on-site early detection tool when coupled with field-based DNA extraction kits. Initially, we optimised the assay for sensitivity and specificity in the lab using gold-standard lab-based DNA extraction; this allowed us to identify required optimization steps more readily at the initial assay development stage. Moreover, the assay was validated with field samples collected around the island of Crete, Greece, which now has established *P. miles* populations. This presumed non-uniform distribution, which should lead to variable amounts of *P. miles* eDNA, was the ideal setting to test our new quantitative assay. Additionally, given the ability of qcLAMP to amplify DNA from crude samples, we evaluated this approach as an alternative qualitative detection tool that can be applied on-site without the need for DNA extraction.

## Materials and methods

### Sample collection

#### Fish tissue samples

For the initial assay development, three *P. miles* individuals were obtained from local fishermen caught off the island of Dia, approximately seven nautical miles north of Heraklion, Crete. Individuals were dissected and 1 cm^2^ muscle tissue samples were obtained and stored at -80 °C until further analysis. Additionally, another four samples of *P. miles* DNA from tissues of individuals collected in Rhodes, were provided by the Hellenic Centre for Marine Research (HCMR) from Heraklion, Crete. To test the assay specificity against closely related species, one individual of each of *Scorpaeana scrofa, Mullus barbatus* and *Mullus surmuletus* were also obtained from local fish markets in Heraklion, Crete. Further specimens of *Scorpaena porcus* and *Scorpaena notata* collected from the Thracian Sea, were provided by the Fisheries Research Institute (INALE), Nea Peramos, Kavala (Hellenic Agricultural Organization-DIMITRA).

#### Aquarium water samples

The performance of the assay on target eDNA from seawater was initially assessed on the following samples: i) water samples were collected from the *P. miles* aquarium tank at CRETAquarium (Heraklion, Crete, Greece); ii) one aquarium sample provided by colleagues at Dublin City University, Ireland; iii) water samples containing closely related and other local species, but lacking *P. miles*, from the fish tanks of the Aquaworld aquarium (Hersonissos, Crete, Greece). For each water sample, 1 L of aquarium tank water was passed through a 0.45 μm pore size polycarbonate filter (Cyclopore, Whatman, Germany) shortly after collection. Filters were stored at -80 °C until further analysis.

#### Field samples

Natural samples were collected from nine coastal sites in Crete, where *P. miles* had been previously sighted (sightings by fishermen, divers, and citizens recorded on gbif.org), during November 2021 and May 2022 (**Figure 1**). Sampling was carried out from either a stand-up paddle board or a small boat using a 2 L Niskin bottle (Aquatic Biotechnology, Spain). Seawater was collected from a depth of 10-20 m, and returned to the beach. Two aliquots of 0.5 L were filtered through Sterivex filter cartridges (HV 0.45 μm, Millipore, USA) using sterilized 60 ml disposable syringes, to provide technical replicates. Additionally, 1 L of bottled deionized water was filtered at each station, to serve as a negative control in the field. Following filtering, filter units were stored in individual ziplock bags on dry ice until returned to the laboratory, where they were stored at -80 °C until further analysis.

**Figure 1.**
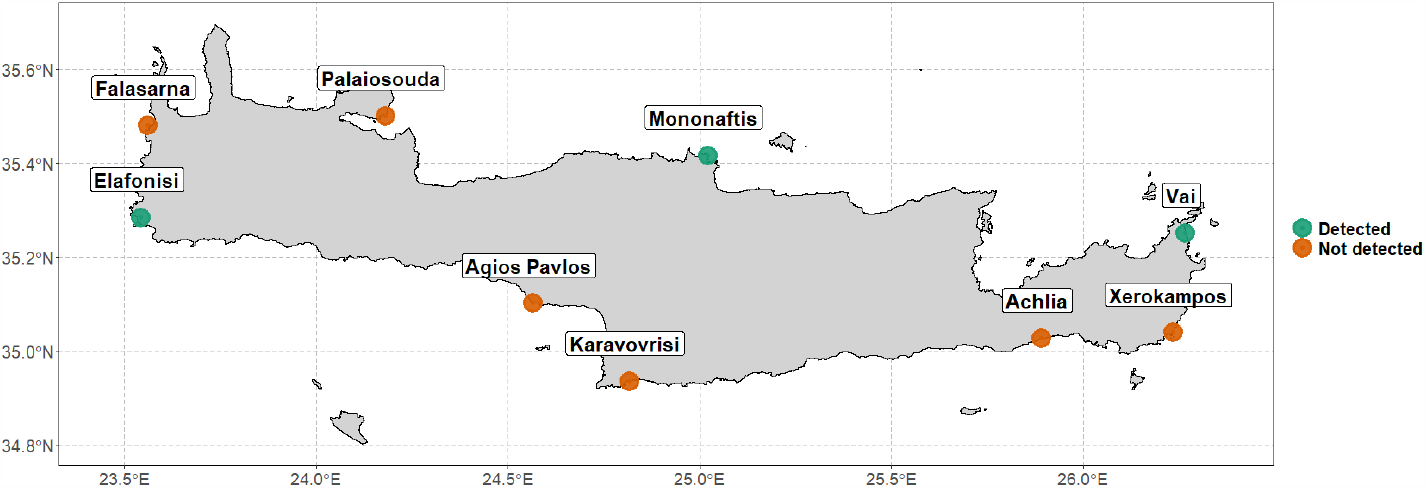
eDNA sampling locations around Crete. Green dots denote that *P. miles* was detected and orange dots denote that it was not detected.

### DNA extraction

#### Tissue samples

Genomic DNA was extracted from tissue samples using the DNeasy Blood & Tissue Kit (Qiagen, Germany). Fish muscle tissue was cut into small pieces and processed following the manufacturer’s instructions with a modified lysis incubation of 3 hours. The quality of extracted DNA was checked on a 1% agarose gel, purity was assessed using a Nanodrop spectrophotometer (ND-1000, Nanodrop Technologies Inc., USA), and DNA concentrations were confirmed with the Qubit 4 (Thermofisher Scientific) using the Qubit 1X dsDNA HS Assay kit (ThermoFisher Scientific).

#### Environmental DNA

Environmental DNA was extracted from Sterivex filters following sterile procedures in a dedicated laboratory, separate from other work involving *P. miles* tissues and amplicons. Sterivex filters were removed from the cartridge and the filter was cut to small pieces (approximately 0.4 cm^2^) using a sterile single-use scalpel and petri dish. eDNA was extracted from filters using the DNeasy Blood & Tissue Kit following the same protocol as for the tissues, with one modification. To ensure the complete disruption of particles containing eDNA, the lysate was passed through a QIAshredder (Qiagen, Germany) column as per the DNeasy plant extraction kit, prior to binding the DNA on the extraction membrane. Single use and sterile consumables were changed between samples (e.g., gloves, scalpel blades, petri dishes).

### Sequencing of COI from reference tissue and aquarium water samples

To guide the assay development and confirm the identity of target and non-target fish, we partially sequenced the mitochondrial cytochrome oxidase subunit 1 (COI) gene. For tissues excised from individual fish, we used the Universal fish PCR primers, FISH-BCL (Forward) and FISHCOIHBC targeting a 704 bp fragment of the COI gene (40). For *P. miles* aquarium water samples, which could contain DNA from other fish in the tank or from the fish food, we designed a species-specific primer; PmCOI7F was used in conjunction with primer FISHCOIHBC and the same PCR conditions to specifically amplify *P. miles* COI from the mixed population (**Table 1**); primer design was based on a 606 bp consensus sequence of 74 GenBank sequences for *P. miles* COI gene. The GenBank sequences were collated in Geneious Prime 2014.7.1.3 and aligned using MAFFT pairwise global alignment (41). Primer PmCOI7F was combined with the universal FISHCOIHBC for species-specific amplification of a 572 bp COI gene fragment.

**Table 1.**
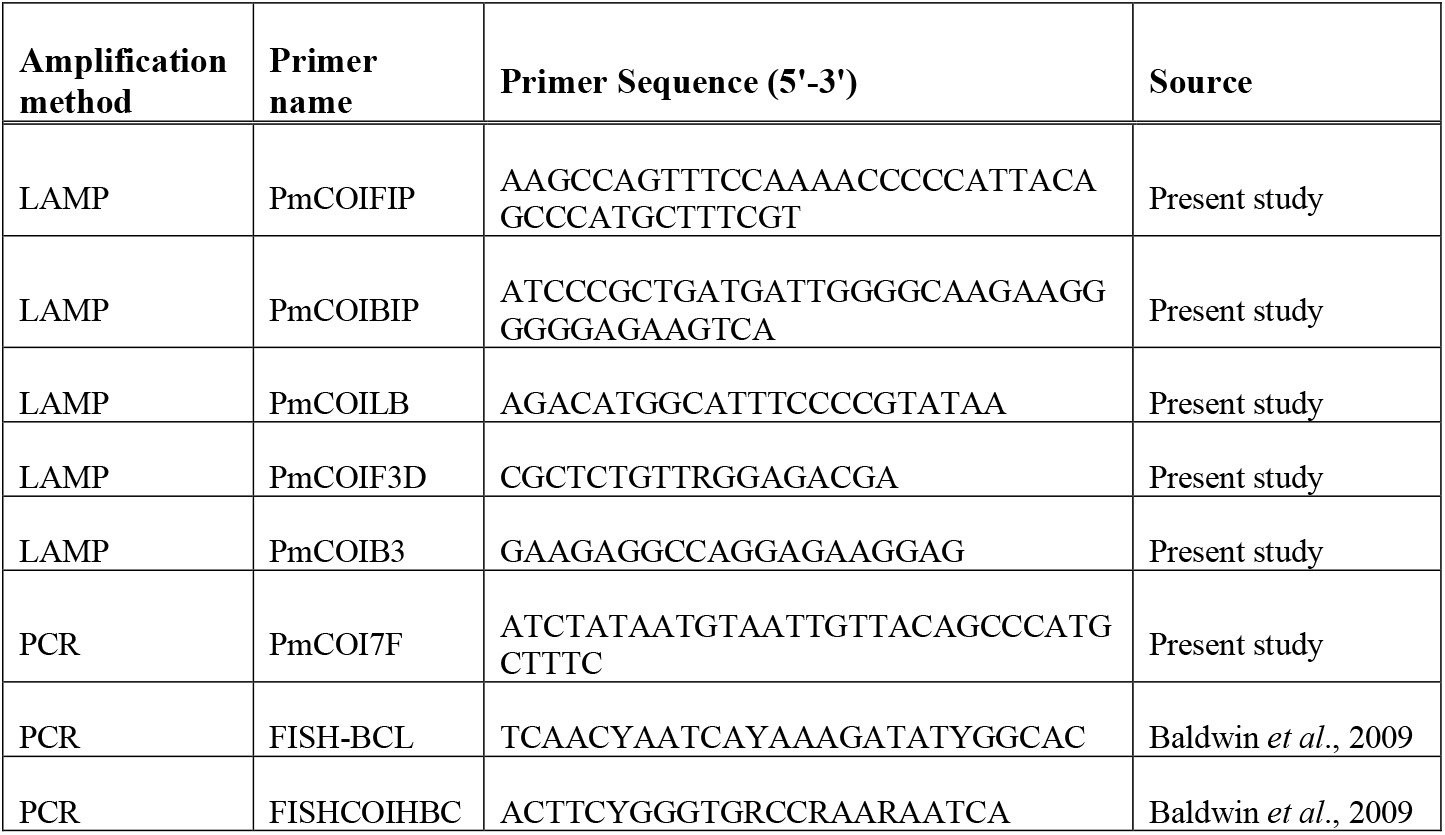
Primers used to amplify and sequence *P. miles* DNA in this study.

PCR on DNA extracted from fish tissues with primers FISH-BCL and FISHCOIHBC) as well as on eDNA extracted from aquarium water samples (with primers PmCOI7F-FISHCOIHBC) was performed in 25 μl reactions which included: 12.5 μl KAPA2G Fast HotStart mix (KAPA Biosystems, UK), 1.25 μl forward primer (10 μM), 1.25 μl reverse primer (10 μM), 9 μl molecular grade water (Sigma-Aldrich), 1 μl template. The PCR cycles were as follows: initial denaturation at 95 °C for 3 min, 35 cycles of denaturation at 95 °C for 15 s, annealing at 60 °C for 30 s and extension at 72 °C for 30 s, and a final extension at 72 °C for 1 min. PCR amplicons were Sanger sequenced by Genewiz (Germany). Forward and reverse reads were paired in Geneious Prime 2014.7.1.3 and species identity was confirmed with the Basic Local Alignment Search Tool (BLAST) on NCBI. Sequences have been submitted to GenBank under accession numbers OR794518 -OR794524.

### qcLAMP assay design

Primer design. Primers were designed based on the *P. miles* COI consensus sequence, produced from sequences obtained in this study as well as from GenBank (**Tables S1 and S3**). Haplotypes were generated using DnaSP 6.12.03 with the default parameters (42). LAMP primers were designed using PrimerExplorer V5 (https://primerexplorer.jp/e/) to target a 213 bp fragment of the *P. miles* consensus sequence for the mitochondrial gene COI (**Figure 2**). This resulted in 5 LAMP primers: Forward inner primer (FIP), backward inner primer (BIP), forward outer primer (F3), backward outer primer (B3) and only a single loop primer, the backward primer (LB). A forward loop primer (LF primer) was not predicted by the software due to no suitable sequence regions. The forward primer F3 was designed with a single degeneracy due to polymorphism at the binding sites. The generated primers were examined *in silico* in pairs for their specificity to the target by using Primer BLAST (NCBI GenBank). Potential cross-reaction and secondary structures in the primers were identified using Primer Digital (https://primerdigital.com/). The final primer sequences are shown in **Table 1**.

**Figure 2.**
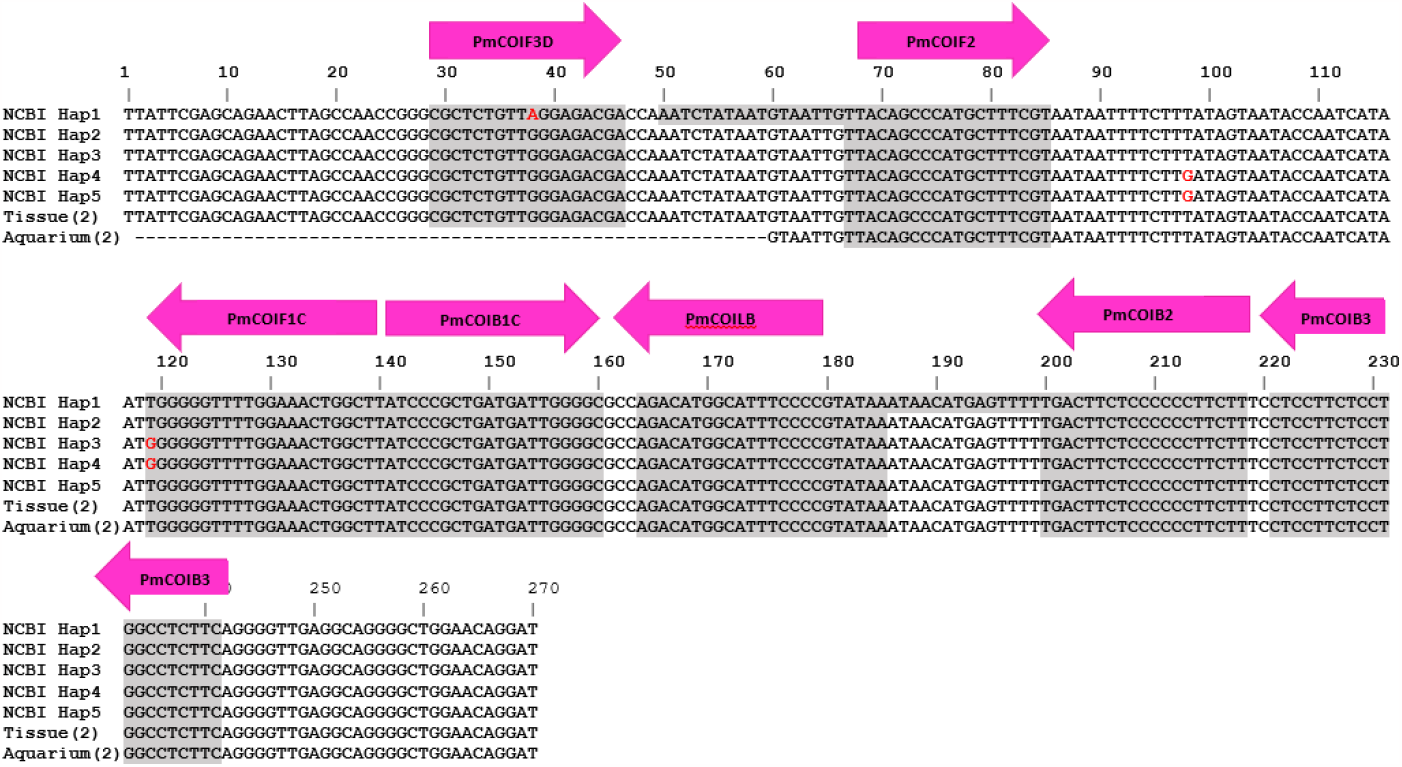
Alignment of the five *P. miles* haplotypes based on GenBank sequences and sequences obtained in this study (Table S1). Hap is short for Haplotypes; the 5 haplotypes shown in the alignment were the ones with sequence variation in the primer target region. Grey areas denote primer binding sites. Arrows show the direction and names of the LAMP primers. The naming of the binding sites follows the standard notation in LAMP primer design since a single primer binds in two positions of the sense and antisense DNA(43).

### Quantitative colorimetric LAMP assay

#### Device

The newly developed qcLAMP device used in this work has been described in Papadakis *et al*. (39). Briefly, it consists of 3D-printed mechanical and electrical parts assembled to a light, hand-held device portable prototype device which is operated by a mobile app. It includes a heating element and a camera to measure colour change within eight reaction-tubes, resulting in a real-time amplification curve based on Colour Index; photo-shots of the camera are collected and transferred via a bluetooth to a nearby tablet or smartphone. For field-based detection, the commercially available version of the device (BIOPIX DNA Technology, GR) was used. The research-based device, available under the brand name “Pebble-R”, employs six reaction tubes and can be used together with a commercial LAMP mix or home-made reaction assays.

#### Assay

Initial optimization for the optimal LAMP incubation temperature was determined using fluorescence real-time LAMP following manufacturer’s instructions (WarmStart LAMP kit, New England Biolabs) and a standard laboratory qPCR thermal cycler (CFX Connect, BIO-RAD) due to higher throughput. The qcLAMP reaction contained 12.5 μl WarmStart Colorimetric LAMP 2X Master Mix with UDG to prevent amplicon contamination (New England Biolabs, USA), 2.5 μl primer mix (FIP 1.8 μM, BIP 1.4 μM, LB 0.6 μM, F3 0.3 μM, B3 0.2 μM), 4 μl DNA template and 6 μl molecular grade water (Sigma-Aldrich), to a final reaction volume of 25 μl. For both tissue and natural seawater sample, 4 μL of the extracted DNA was added to each reaction with a total volume of 25 μL. Reactions were overlayed with 15 μL mineral oil (Sigma-Aldrich) to prevent evaporation and were run in the device at 65 °C for 40 min. Time to positivity (TTP) for each sample was determined by using the background values prior to the onset of amplification, as negative controls could not be included in every run due to the 8 position limit. The threshold colour index was set as the average background value plus 3 times the standard deviation (44) and the TTP was the time it took for the colour index to rise above this value.

### eDNA detection from crude samples

Although the developed assay can be performed on a portable device, as yet, there is no validated eDNA extraction protocol that can be performed in the field. This is a bottleneck for a sample-to-result analysis at the point of sampling. Since LAMP is known to be tolerant to crude samples containing sample inhibitors (45-47), we tested whether *P. miles* could be detected directly from the concentrated seawater particles on the filter, without performing an extraction. We cut small pieces of the filter and added them directly into the colorimetric LAMP reactions and performed a heat lysis step at 90 °C for 9 min on the same device prior to amplification. We first confirmed that a heat lysis incubation did not affect the functionality of the LAMP reagents, using *P. miles* tissue extracted as positive controls. Filter pieces from the *P. miles* aquarium tank samples were then used as simulated environmental samples. Small square filter pieces approximately 0.4 x 0.4 cm were cut from the total 4 x 4 cm filter, and a single piece was added directly to 25 μL colorimetric LAMP reactions. Following the lysis incubation, reactions were cooled down for 5 min, and re-introduced to the device for amplification to commence.

## Results

In this study, we initially designed a species-specific LAMP assay for *P. miles*, based on sequences from NCBI and from locally collected fish samples. We established that the assay is specific to the target species using both tissue samples and eDNA mock samples collected from aquarium tanks, with the target species present or absent. Finally, we tested the assay on field samples collected around Crete, Greece. In addition, we tested whether we could detect *P. miles* from crude eDNA samples without a DNA extraction step so that the whole analysis could be performed in the field. The overall approach is depicted in **Figure 3**. In workflow A, extracted DNA from either tissue or seawater samples was added to the qcLAMP reactions, and in workflow B, pieces of the filter were added directly without extraction of the captured particles. Target amplification in the device was monitored in real time, generating amplification curves where the faster the signal rises above the background, the more of the target gene is present.

**Figure 3.**
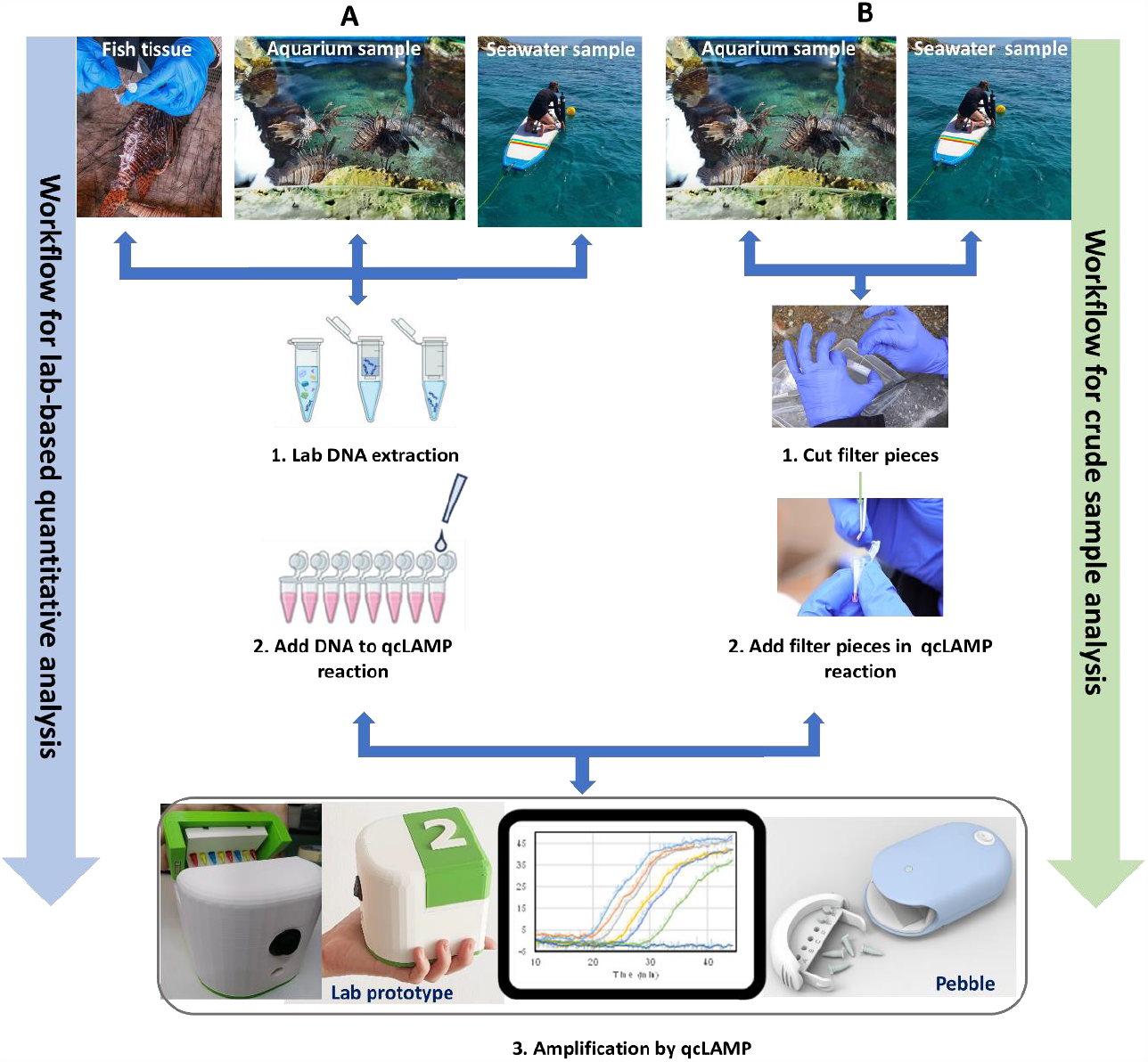
Assay development process used in this study. Increasing complexity of sample type, from fish tissue to aquarium and field (seawater) samples. (A). Testing the quantitative capacity of the assay with DNA from all three samples extracted in the laboratory, as well as (B) the capacity to detect the target from small pieces of the filter in the last two cases of samples (aquarium and sea).

### Quantitative colorimetric LAMP assay optimization

The new LAMP assay was designed based on *P. miles* sequences obtained from NCBI and locally. Tissues from local fish revealed two haplotypes, DNA from the *P. miles* tank water of the CRETAquarium matched one of these haplotypes, and a different third haplotype was obtained from *P. miles* tank water of the Dublin Aquarium (**Figure 2**). The assay was initially evaluated using DNA extracts from fish tissues. Primer concentrations were optimized for reaction speed and to reduce non-specific amplification. The forward primer F3 contained a two-fold degeneracy i.e., two different primer sequences at half the concentration each. Therefore, increasing the concentration of this primer by 50% relative to the standard protocol, resulted in a faster onset of amplification. Decreasing the concentration of the BIP primer resolved occasional non-specific products in the negative controls, as *in silico* analysis had predicted a cross-reaction between the primers BIP and LB. To assess if a second loop primer would improve the reaction speed, a forward loop primer (LF) was manually designed and tested. There was no difference in reaction speed with the inclusion of LF, therefore, this primer was omitted. All the above adjustments were adopted in our qcLAMP assay and subsequent experimentations.

### qcLAMP assay sensitivity and specificity

The assay’s sensitivity was determined by serially diluting and amplifying *P. miles* tissue extracted DNA with LAMP. This test established the quantitative capacity of the assay, the LoD and the reproducibility among replicates. For initial tests, we used real-time LAMP with a fluorescence dye to obtain an assay benchmarking from an established laboratory device (i.e., the qPCR thermal cycler) prior to proceeding with colorimetric LAMP on the prototype colorimetric device. To enable direct comparisons, both tests (fluorescent LAMP and qcLAMP) were performed in parallel on the same DNA samples. These were a gDNA dilution series at the following concentrations: 20 ng, 2 ng, 0.2 ng, 0.02 ng, 0.002 ng, in triplicate reactions. The assay performed linearly across the whole range tested in both fluorescence and qcLAMP (**Figure 4**). The LoD was determined as the lowest DNA concentration at which 95% of replicates produced a positive result (48). To ensure that the LoD was comparable to other reported studies, we estimated the number of COI copies in *P. miles* genomic DNA by qPCR (LoD at 0.002 ng is equivalent to 50 gene copies; **Figure S1, Table S2**). qcLAMP generally showed slower reaction times than fluorescence LAMP presumably due to device differences. At the LoD of 0.002 ng μL^-1^ reaction, the TTP for qcLAMP was 31.2 min, while for fluorescence LAMP it was 25.1 min, with high variability among replicates.

**Figure 4.**
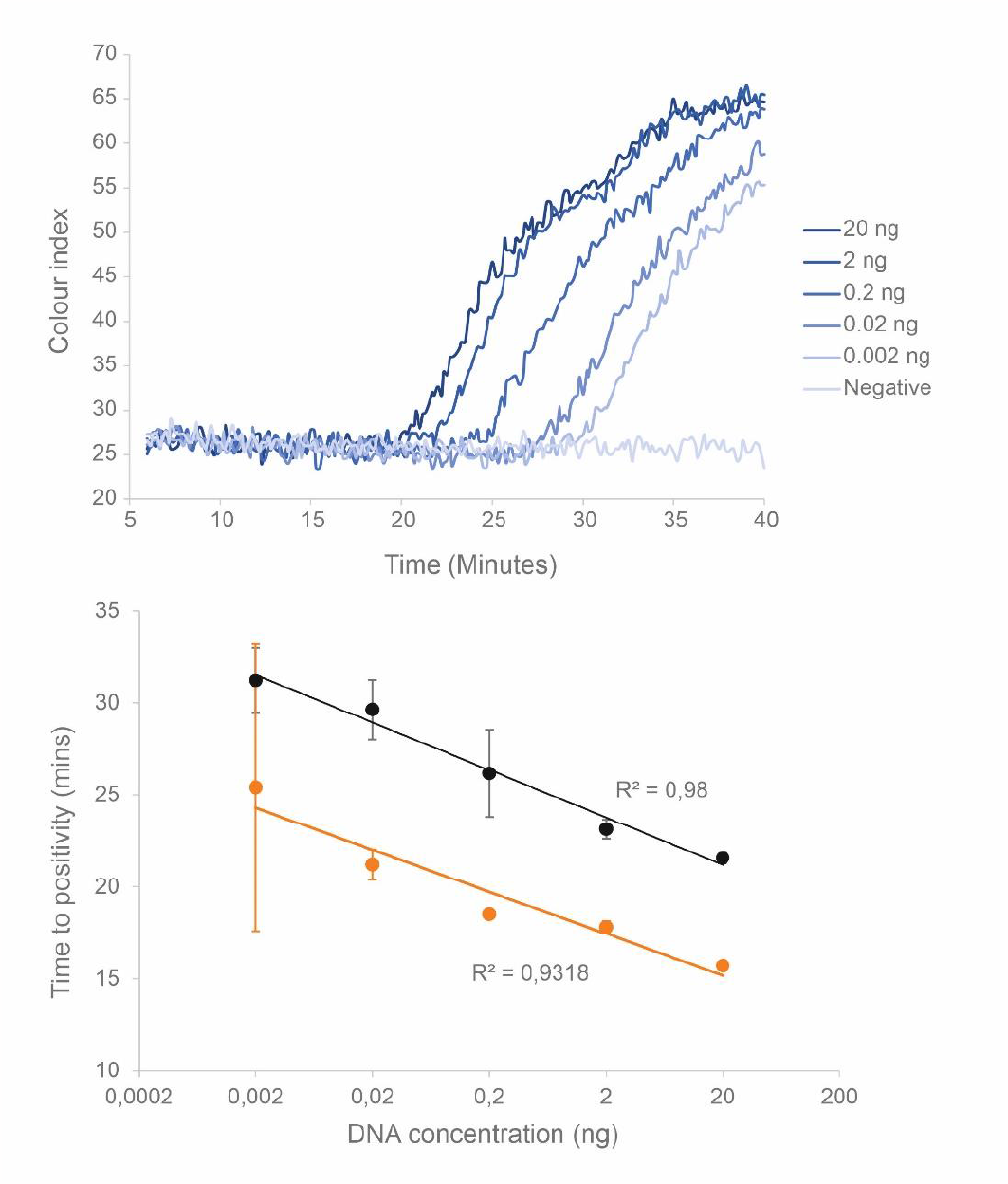
Sensitivity and quantitative range of the real-time *P. miles* LAMP assay established by serial dilutions of genomics DNA extracted from fish tissues (n=3). (Top) Real-time amplification curves for the different DNA concentrations, showing later onset of amplification for lower concentrations. (Bottom) Linear relationship of the corresponding TTP values for qcLAMP (black), as well as the equivalent experiment performed using fluorescence LAMP on a qPCR cycler (orange).

The assay specificity was evaluated by testing against species closely related to the target as well as co-occurring species in the area. Samples were obtained both from fish tissue and from aquarium tanks containing *P. miles* and others that did not contain *P. miles*, simulating environmental negatives (**Table 2**). Species of the *Scorpaena* genus were prioritised in the tests as these are members for the Scorpenidae family and the closest relatives of *P. miles* in terms of sequence identity of the COI gene (*S. porcus*: 81.1%, *S. scrofa*: 82.3%, and *S. notata*: 81.4%). Specificity was further tested against fish species that typically co-occur with the *P. miles* in eastern Mediterranean waters. In total, 18 non-target species and six *P. miles* individuals were tested for primer specificity. There was no amplification from DNA of species closely related to *P. miles*, or DNA from aquarium tanks containing non-target species (**Table 2**), confirming that the assay is species specific.

**Table 2.**
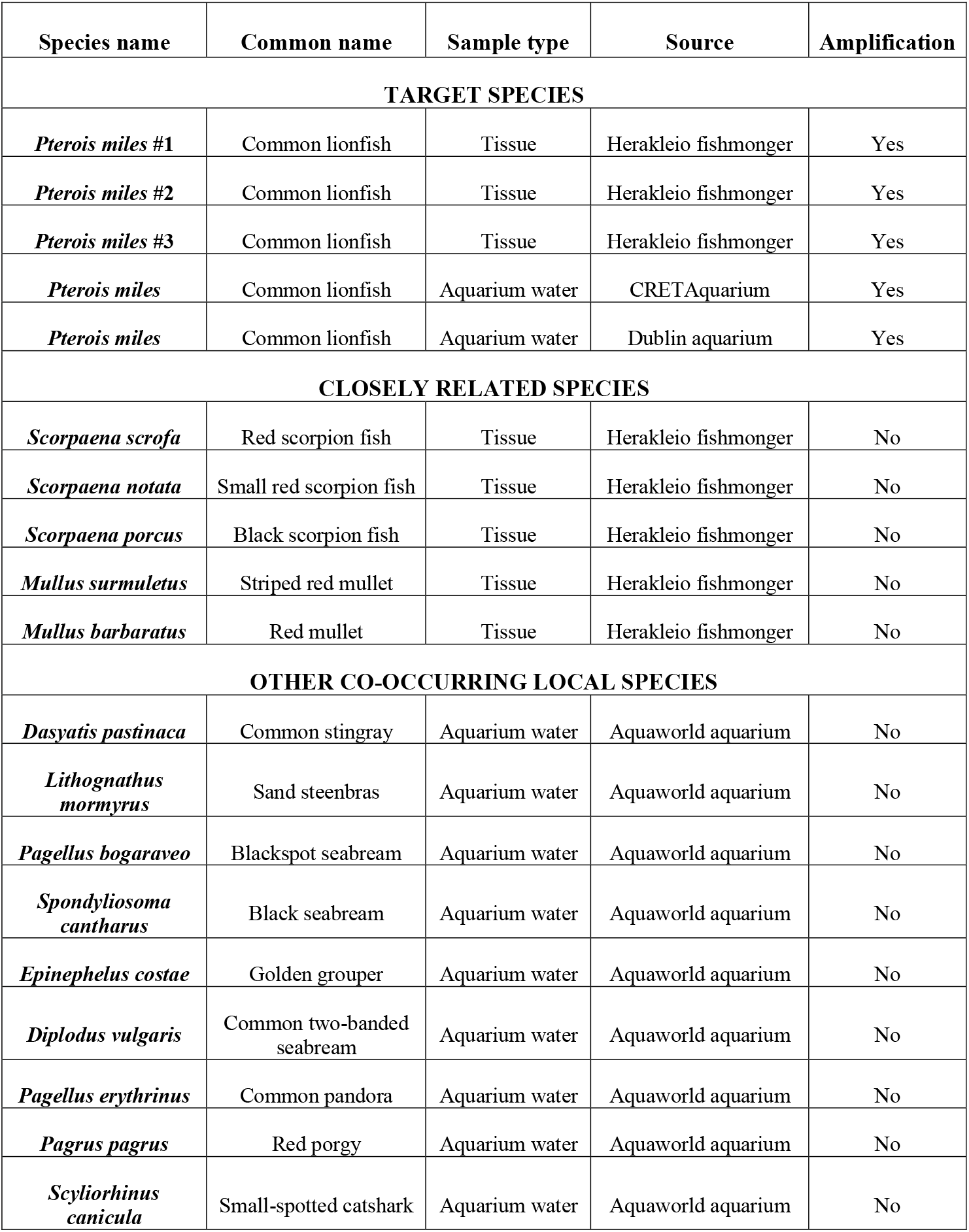
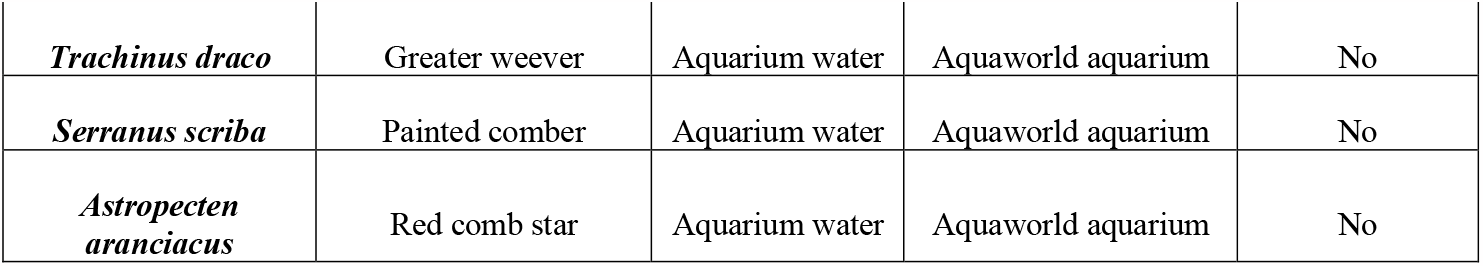
Results of assay specificity testing showing fish species tested, including sample type and source. Three individuals of *P. miles* from which tissue was obtained are marked with a #.

### Assay validation using natural and aquarium eDNA samples

The assay was validated by testing the *P. miles* qcLAMP protocol on seawater samples collected from nine locations around the island of Crete, Greece (**Figure 1**) as well as samples collected from the CRETAquarium and Dublin aquarium tanks. Following eDNA extraction, the amplification quality of the DNA was confirmed spectrophotometrically, fluorometrically and by PCR using the Universal fish primers (**Table S3**). All environmental samples and negative field controls (e.g., bottled water) were analyzed for *P. miles* by qcLAMP using the device. *P. miles* eDNA was detected in three of our nine field samples (**Figure 1, Table 3**); North Central Crete (Mononaftis beach), Eastern Crete (Vai beach), and Western Crete (Elafonisi beach), with the latter two calculated as having a considerably higher DNA concentration than the former. The target concentration in positive reactions was determined using the DNA standard curve equation. To extrapolate the DNA concentration per reaction, to the DNA concentration in the original sample in units of ng L^-1^, we accounted for the volume of seawater filtered per sample (**Table 3**), the initial elution volume of the DNA extraction (50-100 μL), and the amount of DNA added per reaction (4 μL). We calculated *P. miles* DNA concentrations in natural seawater ranging from ∼ 0.15 – 3.19 ng L^-1^ and in the two aquarium samples 2.86 ng L^-1^ and 3.91 ng L^-1^ (**Table 3**). These concentrations are in the middle of the tested assay range.

**Table 3.** Details of sample volumes, TTP and calculated concentrations of eDNA recovered from natural and aquarium seawater samples.

| Sampling location | Seawater volume filtered (L) | Eluted volume of DNA (µL) | Time to Positivity (mins) | Calculated *P. miles* DNA concentration in original sample (ng L<sup>-1</sup>) |
| --- | --- | --- | --- | --- |
| Elafonisi | 1 | 80 | 26.6 | 3.19 |
| Vai | 1.56 | 50 | 25.6 | 3.12 |
| Mononaftis | 1 | 50 | 29.5 | 0.15 |
| CRETAquarium | 0.4 | 70 | 27.6 | 2.86 |
| Dublin Aquarium | 0.5 | 100 | 27.4 | 3.91 |

### Direct analysis of crude seawater samples

Amplification of *P. miles* eDNA was feasible directly from the filter used to collect eDNA containing particles from the seawater. This was achieved with a simple heat-lysis step performed within the qcLAMP device immediately prior to amplification. We initially applied this protocol to samples collected from the CRETAquarium *P. miles* tank. A positive amplification was achieved with a TTP of 30 min, equivalent to 0.008 ng DNA. Extrapolating the amount of DNA from the filter piece to the equivalent for the whole filter membrane, and accounting for the volume filtered, the sample quantifications yielded a value of 2.5 ng L^-1^, which is similar to the 2.86 ng L^-1^ obtained previously for the same tank using extracted DNA samples (**Table 3**).

## Discussion

### LAMP optimization for target detection in eDNA samples

The scope of this study was to develop a simple and fast molecular assay that can accurately quantify the invasive *P. miles* from seawater samples, which is also applicable in the field. The LAMP method was selected since it is ideal for species-specific assays due to the six primers required along with the efficient production of amplicon. This provides the sensitivity and specificity to eliminate the risk of false positives from non-target amplification, and false negatives from low sensitivity (42). However, several optimization steps were required to guarantee the reliability of the *P. miles* qcLAMP assay. An inherent limitation and risk of the LAMP method is the formation of primer dimers due to the high number of primers used (34). We overcame this issue by recognising these interactions *in silico* and optimizing primer concentrations. We confirmed our assay’s ability to amplify different *P. miles* haplotypes to ensure its potential application on a regional scale. We also demonstrated the specificity of our assay in the presence of non-target species by testing against species of the same family (Scorpaenidae) and against additional other species that naturally occur in the regions that we conducted our field analyses. In addition to fish tissue samples, we used water samples from aquarium tanks that contained multiple local marine species to simulate a natural environmental sample. Our assay showed a reliable and highly sensitive detection as low as 0.002 ng per reaction of the mitochondrial COI gene, comparable to other studies using LAMP as their detection method for eDNA in aquatic environments (49-51). This is approximately equivalent to 25 gene copies per reaction, which is within the usual LoD for LAMP and other quantitative DNA amplification methods, such as qPCR and qRPA. Interestingly, we noted that at this low DNA input, qcLAMP on the prototype device outperformed the standard fluorescence real-time LAMP performed on the qPCR cycler. This might be due to differences in the operation of the two devices, or the presence of inhibitors that may affect fluorophores.

### Quantification of *P. miles* eDNA in field samples

Following assay design, optimization, and *in vitro* testing, the protocol was assessed on eDNA from natural samples collected from nine locations around Crete. The target species was detected in three out of the nine locations (**Figure 3**). These three locations are either semi-enclosed bays (Mononaftis) which are sheltered from onshore winds and waves, or are not exposed to prevailing winds throughout the year (Vai and Elafonissi). The highest eDNA concentrations of 3.12 ng L^-1^ and 3.19 ng L^-1^ were found at Vai and Elafonissi beaches, which were similar to those found in the aquarium water tanks. Even though these areas were sampled in different seasons (November 2021 and May 2022, respectively), both are protected nature reserves where fishing is not allowed and are therefore expected to have a higher overall fish biomass. Similar findings have been reported in Cyprus, where the highest densities of *P. miles* populations were recorded in Marine Protected Areas where fishing is forbidden (16). Mononaftis bay, which was sampled in May 2022, contained a lower amount of eDNA by one order of magnitude, even though it is SCUBA diving location due to it being rich in marine life. However, at the time of sampling the waters were still at pre-summer temperatures and only two individuals were observed by divers at the sampling location.

The lack of detection in the remaining regions could be attributed to potentially lower population densities of the target species, as well as to various biotic and abiotic factors that could affect eDNA concentration, distribution and persistence in the water column (20). The effectiveness of eDNA detection depends on its persistence in the water column at the sampling point, which can, in turn, depend on its release from the target organism, temperature of water, transport across space and decay rates (32, 52-54). Although eDNA is concentrated by filtering water samples, the relative concentration of target DNA can remain very small, thus making its detection particularly challenging (29). In the case of our target, its ecology and habitat preferences play a major role in the presence and distribution of its extra-organismal DNA. *P. miles* static lifestyle could limit the distribution of eDNA in the water column in calm waters without currents. Moreover, this species migrates to deeper waters in winter, likely to shelter from rougher weather conditions at the surface (55). A solution could be to filter larger volumes of water instead of the 1 L filtered in this study and perform more frequent seasonal sampling.

### On-site *P. miles* qcLAMP eDNA detection directly from the sample

A major bottleneck for field-based eDNA molecular assays is DNA extraction. LAMP has been demonstrated to provide sensitive target amplification from a number of crude sample matrices, including food and plant tissues (47, 56, 57). We tested whether we could amplify and detect *P. miles* eDNA directly from the filter that contained concentrated biomass, including tissue particles from *P. miles*. We successfully released DNA for amplification by using a portable, commercially available qcLAMP device and by directly inserting filter pieces in the LAMP reaction and preheating the reaction at 90 °C. Comparison to equivalent samples that were amplified after DNA extractions showed that LAMP was unaffected by remaining lysed particles. This agrees with previous results with LAMP using crude samples, where protozoan and bacterial DNA was amplified without a DNA extraction (39, 58, 59). This approach is useful for on-site detection, although using only 1 % of the total filter for amplification obviously reduces the overall sensitivity for field samples. Further testing is needed at locations with visually evident populations to establish the required LoD for management purposes.

## Conclusion

In conclusion, we have developed and validated a fast, sensitive and species-specific qcLAMP assay combined with a novel prototype and commercially available qcLAMP device to detect and quantify eDNA of the marine invasive lionfish, *P. miles*. The assay was fully optimized for sensitivity, specificity, and quantitative capacity. We demonstrated the assay using the rapid and simple-to-use portable device which can be taken to the field for on-site analysis. We also demonstrated that on-site *P. miles* detection is feasible by direct amplification from the sample.

We have used our assay to detect the presence of *P. miles* in natural areas around Crete, where the species has been previously sighted. The presence of *P. miles* was detected in three out of nine locations: these are nature reserves protected from fishing and boating, and may thus have higher population densities of the target species. Our results highlight the need for further standardizing the method in the field, with targeted studies of eDNA concentration and dispersal patterns in relation to species life history and hydrographic conditions so that quantitative eDNA results can be related to actual population distributions. Once fully standardized for application in the field, this will be and easy-to-use monitoring tool for the early detection of *P. miles* invasion routes, for collecting information about this species’ dispersal patterns and for providing timely data to policy makers to manage *P. miles* invasions. Our proposed qcLAMP methodology is generic and widely applicable for other species in marine and fresh waters, thus presenting a new tool for ecological monitoring.

## Acknowledgments

Dr. Aspasia Sterioti for access to the aquarium tanks at CRETAquarium, John McLaren for access to the aquarium tanks at Aquaworld Hersonissos, Philipos Marakis and Dr. Aggelos Ioannou at the Ecodive Center Heraklion for help with sampling at Mononaftis Bay and visual confirmation of lionfish presence, and Dr. Fiona Bracken at Dublin City University for aquarium samples.

## Author contributions

KHM, MV, PK, CG and EG designed the study and experiments. KHM, MV, CG, PK and PX performed sample collections and experiments. KHM, MV and EG wrote the manuscript. All authors contributed to the manuscript and approved the submitted version.

