## Supplementary material for "Development of a quantitative colorimetric LAMP assay for fast and targeted molecular detection of the invasive lionfish *Pterois miles* from environmental DNA": Supplemetary information

**SUPPLEMENTARY INFORMATION**

**Sequences used for assay design**

A total of 74 sequences were retrieved from GenBank in March 2021. Whole COI sequences were initially grouped into Haplotypes and consensus sequences of each group were used for the primer design. For illustration purposes, Figure 2 in the main text shows Haplotypes that differed only in the LAMP amplicon region, including the primer binding sites. These were renamed sequentially for the purposes of the Figure.

Table S1. Information for the sequences used to design the LAMP assay.

| Accession no | Location | Authors | DOI | Haplotype no. based on whole sequence | Haplotype no. based on amplicon region only (as shown in Fig. 2) |
|---|---|---|---|---|---|
| MF124021 | Israel | Kimmerling et al., 2018 | https://doi.org/10.1038/s41559-017-0413-2 | Haplotype A | Haplotype 1 |
| MN150238 | Cyprus | Dimitriou et al., 2019 | https://doi.org/10.3390/d11090149 | Haplotype B | Haplotype 2 |
| MN150237 | Cyprus | Dimitriou et al., 2019 | https://doi.org/10.3390/d11090149 | | |
| MN150236 | Cyprus | Dimitriou et al., 2019 | https://doi.org/10.3390/d11090149 | | |
| MN150235 | Cyprus | Dimitriou et al., 2019 | https://doi.org/10.3390/d11090149 | | |
| MN150234 | Cyprus | Dimitriou et al., 2019 | https://doi.org/10.3390/d11090149 | | |

| | | | |
|---|---|---|---|
| MN150233 | Cyprus | Dimitriou et al., 2019 | https://doi.org/10.3390/d11090149 |
| MN150232 | Cyprus | Dimitriou et al., 2019 | https://doi.org/10.3390/d11090149 |
| MN150231 | Cyprus | Dimitriou et al., 2019 | https://doi.org/10.3390/d11090149 |
| MN150230 | Cyprus | Dimitriou et al., 2019 | https://doi.org/10.3390/d11090149 |
| MN150228 | Cyprus | Dimitriou et al., 2019 | https://doi.org/10.3390/d11090149 |
| MN150223 | Cyprus | Dimitriou et al., 2019 | https://doi.org/10.3390/d11090149 |
| MN150222 | Cyprus | Dimitriou et al., 2019 | https://doi.org/10.3390/d11090149 |
| MN150221 | Cyprus | Dimitriou et al., 2019 | https://doi.org/10.3390/d11090149 |
| MN150220 | Cyprus | Dimitriou et al., 2019 | https://doi.org/10.3390/d11090149 |
| MN150219 | Cyprus | Dimitriou et al., 2019 | https://doi.org/10.3390/d11090149 |
| MN150218 | Cyprus | Dimitriou et al., 2019 | https://doi.org/10.3390/d11090149 |
| MN150214 | Cyprus | Dimitriou et al., 2019 | https://doi.org/10.3390/d11090149 |
| MN150211 | Cyprus | Dimitriou et al., 2019 | https://doi.org/10.3390/d11090149 |
| MN150210 | Cyprus | Dimitriou et al., 2019 | https://doi.org/10.3390/d11090149 |
| MN150209 | Cyprus | Dimitriou et al., 2019 | https://doi.org/10.3390/d11090149 |

| | | | |
|---|---|---|---|
| MN150208 | Cyprus | Dimitriou et al., 2019 | https://doi.org/10.3390/d11090149 |
| MN150206 | Cyprus | Dimitriou et al., 2019 | https://doi.org/10.3390/d11090149 |
| MN150204 | Cyprus | Dimitriou et al., 2019 | https://doi.org/10.3390/d11090149 |
| MN150201 | Cyprus | Dimitriou et al., 2019 | https://doi.org/10.3390/d11090149 |
| MN150200 | Cyprus | Dimitriou et al., 2019 | https://doi.org/10.3390/d11090149 |
| MN150196 | Cyprus | Dimitriou et al., 2019 | https://doi.org/10.3390/d11090149 |
| MN150195 | Cyprus | Dimitriou et al., 2019 | https://doi.org/10.3390/d11090149 |
| MN150192 | Cyprus | Dimitriou et al., 2019 | https://doi.org/10.3390/d11090149 |
| MN150191 | Cyprus | Dimitriou et al., 2019 | https://doi.org/10.3390/d11090149 |
| MN150190 | Cyprus | Dimitriou et al., 2019 | https://doi.org/10.3390/d11090149 |
| MN150188 | Cyprus | Dimitriou et al., 2019 | https://doi.org/10.3390/d11090149 |
| MN150186 | Cyprus | Dimitriou et al., 2019 | https://doi.org/10.3390/d11090149 |
| MN150185 | Cyprus | Dimitriou et al., 2019 | https://doi.org/10.3390/d11090149 |
| KU317873 | Saudi Arabia | Rabaoui et al., 2018 | Unpublished |
| MF124020 | Israel | Kimmerling et al., 2018 | https://doi.org/10.1038/s41559-017-0413-2 |
| JQ350297 | Madagascar | Hubert et al., 2012 | https://doi.org/10.1371/journal.pone.0028987 |

| | | | | | |
|---|---|---|---|---|---|
| JQ350295 | Madagascar | Hubert et al., 2012 | https://doi.org/10.1371/journal.pone.0028987 | | |
| JQ350296 | Madagascar | Hubert et al., 2012 | https://doi.org/10.1371/journal.pone.0028987 | | |
| JF494332 | South Africa | Steinke et al., 2016 | https://doi.org/10.1139/gen-2015-0212 | | |
| MT881566 | Indonesia | Nuryanto et al., 2021 | https://doi.org/10.1051/e3sconf/202132201004 | | |
| MT881563 | Indonesia | Nuryanto et al., 2021 | https://doi.org/10.1051/e3sconf/202132201004 | | |
| EU148592 | India | Lakra et al., 2010 | https://doi.org/10.1111/j.1755-0998.2010.02894.x | | |
| EU148593 | India | Lakra et al., 2010 | https://doi.org/10.1111/j.1755-0998.2010.02894.x | | |
| FJ584034 | Indonesia | Steinke et al., 2009 | https://doi.org/10.1371/journal.pone.0006300 | | |
| FJ584033 | Indonesia | Steinke et al., 2009 | https://doi.org/10.1371/journal.pone.0006300 | | |
| FJ584032 | Indonesia | Steinke et al., 2009 | https://doi.org/10.1371/journal.pone.0006300 | | |
| FJ584030 | Sri Lanka | Steinke et al., 2009 | https://doi.org/10.1371/journal.pone.0006300 | | |
| FJ584029 | Sri Lanka | Steinke et al., 2009 | https://doi.org/10.1371/journal.pone.0006300 | | |
| FJ584027 | Sri Lanka | Steinke et al., 2009 | https://doi.org/10.1371/journal.pone.0006300 | | |
| JF494333 | South Africa | Steinke et al., 2016 | https://doi.org/10.1139/gen-2015-0212 | Haplotype C | Haplotype 2 |
| GU805078 | South Africa | Steinke et al., 2016 | https://doi.org/10.1139/gen-2015-0212 | Haplotype D | Haplotype 2 |
| MN150229 | Cyprus | Dimitriou et al., 2019 | https://doi.org/10.3390/d11090149 | Haplotype E | Haplotype 3 |
| MN150227 | Cyprus | Dimitriou et al., 2019 | https://doi.org/10.3390/d11090149 | Haplotype F | Haplotype 4 |
| MN150216 | Cyprus | Dimitriou et al., 2019 | https://doi.org/10.3390/d11090149 | | |

| | | | | | |
|---|---|---|---|---|---|
| MN150215 | Cyprus | Dimitriou et al., 2019 | https://doi.org/10.3390/d11090149 | | |
| MN150205 | Cyprus | Dimitriou et al., 2019 | https://doi.org/10.3390/d11090149 | | |
| MN150202 | Cyprus | Dimitriou et al., 2019 | https://doi.org/10.3390/d11090149 | | |
| MN150198 | Cyprus | Dimitriou et al., 2019 | https://doi.org/10.3390/d11090149 | | |
| MN150187 | Cyprus | Dimitriou et al., 2019 | https://doi.org/10.3390/d11090149 | | |
| MN150226 | Cyprus | Dimitriou et al., 2019 | https://doi.org/10.3390/d11090149 | Haplotype G | Haplotype 2 |
| MN150225 | Cyprus | Dimitriou et al., 2019 | https://doi.org/10.3390/d11090149 | | |
| MN150224 | Cyprus | Dimitriou et al., 2019 | https://doi.org/10.3390/d11090149 | | |
| MN150217 | Cyprus | Dimitriou et al., 2019 | https://doi.org/10.3390/d11090149 | | |
| MN150213 | Cyprus | Dimitriou et al., 2019 | https://doi.org/10.3390/d11090149 | | |
| MN150203 | Cyprus | Dimitriou et al., 2019 | https://doi.org/10.3390/d11090149 | | |
| MN150199 | Cyprus | Dimitriou et al., 2019 | https://doi.org/10.3390/d11090149 | | |
| MN150197 | Cyprus | Dimitriou et al., 2019 | https://doi.org/10.3390/d11090149 | | |
| MN150194 | Cyprus | Dimitriou et al., 2019 | https://doi.org/10.3390/d11090149 | | |
| MN150189 | Cyprus | Dimitriou et al., 2019 | https://doi.org/10.3390/d11090149 | | |

| | | | | | |
|---|---|---|---|---|---|
| MN150212 | Cyprus | Dimitriou et al., 2019 | https://doi.org/10.3390/d11090149 | Haplotype H | Haplotype 3 |
| MN150193 | Cyprus | Dimitriou et al., 2019 | https://doi.org/10.3390/d11090149 | Haplotype I | Haplotype 5 |
| MK041049 | Indonesia | Nuryanto et al., 2021 | https://doi.org/10.1051/e3sconf/202132201004 | Haplotype J | Haplotype 2 |
| MT076824 | Arabian Gulf | Ludt et al., 2020 | https://doi.org/10.5281/zenodo.3934741 | Haplotype K | Haplotype 2 |

**Estimation of COI copy number within *P. miles* genomic DNA.**

Development of a quantitative LAMP assay required a dilution series of *P. miles* DNA to generate a standard curve. Whilst adding known amounts of lionfish gDNA is the standard approach, we also sought to obtain this information in copy numbers to ensure the new LAMP assay had a sensitivity comparable to other assays reported in the literature. We, therefore, generated PCR products to use as standards using the FISH-BCL / FISHCOIHBC PCR primer combination. PCR amplicons were checked for specificity by gel electrophoresis, purified using a DNA Purification Kit (Macherey-Nagel, Germany) and accurately quantified using a Qubit 4 instrument (ThermoFisher Scientific) and Qubit dsDNA HS Assay kit (ThermoFisher Scientific). Quantitative PCR (qPCR) was used to determine the copy number of COI in lionfish genomic DNA (gDNA). This was achieved by comparing gDNA serial dilutions to PCR amplicon standards of known copy numbers. We designed new qPCR primers for *P. miles*, PmCOI3F (5' – AATCTATAATGTAATTGTTACAGCCCATGC – 3') & PmCOI3R (5' – GTCAAAAACTCATGTTATTTATACGGGGAA - 3'), targeting a 155bp region of the species' COI gene, proximal to the region used as a target for LAMP. In brief, the 20µl qPCR reaction consisted of 10µl PowerTrack SYBR Green Master Mix (Applied Biosystems, Lithuania), 1µl of 10µM PmCOI3F, 1µl of 10µM PmCOI3R, 7µl molecular grade water (Sigma-Aldrich), and 1µl DNA template. The qPCR cycling modes were: initial denaturation for 95°C for 2 minutes, and 40 cycles of a) a denaturation step of 95°C for 5 minutes, and b) an annealing step of 64°C for 30 seconds. This approach was used to quantify the number of COI gene copies in *P. miles* genomic DNA (Figure S1).

Fig S1. qPCR standard curve using a dilution curve as quantified PCR products as standards, and a gDNA sample dilution series as unknowns

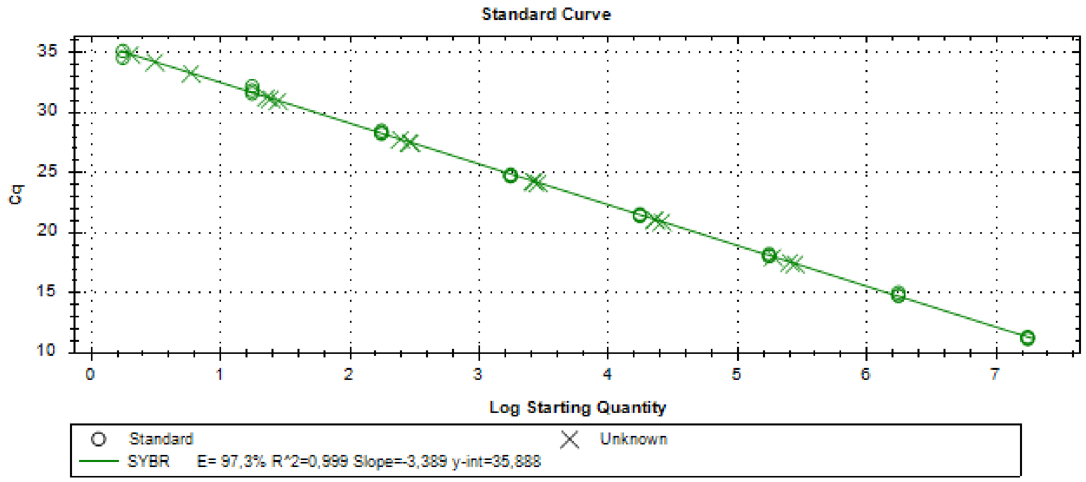

Table S2. Conversion of gDNA standards used for eDNA quantification from field samples, to COI copy number.

| gDNA standard amount (ng) | Copy number |
|---|---|
| 10 | 240297 |
| 1 | 23989 |
| 0.1 | 2718 |
| 0.01 | 276 |
| 0.001 | 25 |
| 0.0001 | 3.7 |

### Field samples collected during the study

Table S3. Field samples collected and results of performed analyses. Where sample volume was limiting (e.g. due to usage in optimization) only qcLAMP was performed. Asterisk indicates that the PCR product was sequenced. SUP stands for Stand-up Paddleboard.

| Location | Latitude | Longitude | Collection date | Collected by | Concentration by Qubit (ng/µl) | Concentration by Nanodrop (ng/µl) | qcLAMP - *P. miles* | qPCR - *P. miles* | PCR – All fish |
|---|---|---|---|---|---|---|---|---|---|
| CretAquarium | - | - | September 2021 | - | 2.5 | 6.5 | Yes | Yes | Yes* |
| Dublin aquarium | - | - | September 2021 | - | 4.5 | - | Yes | Yes | Yes* |
| Vai | 35.25194 | 26.26710 | November 2021 | SUP | 3.14 | 17.9 | Yes | Yes | - |
| Mononaftis | 35.41722 | 25.02006 | November 2021 | Diving School | 1.962 | 9.3 | Yes | Yes | - |
| Falasarna | 35.48143 | 23.55907 | May 2022 | SUP | 1.74 | 11.2 | No | No | Yes |
| Palaiosouda | 35.50157 | 24.17971 | May 2022 | SUP | 2.72 | 7.9 | No | No | Yes |
| Achlia | 35.02783 | 25.89057 | May 2022 | SUP | 2.42 | 7.6 | No | No | Yes |
| Xerokampos | 35.04011 | 26.23361 | May 2022 | SUP | 3.52 | 8.8 | No | No | Yes |
| Agios Pavlos | 35.10223 | 24.56455 | May 2022 | SUP | 2.86 | 10.4 | No | No | Yes |
| Karovovrisi | 34.93637 | 24.81580 | May 2022 | SUP | 3.14 | 9.3 | No | No | Yes |
| Elafonisi | 35.26816 | 23.5261 | May 2022 | Boat | - | - | Yes | - | - |
